# Neuromolecular interactions guiding homeostatic mechanisms underlying healthy ageing: A view from computational microscope

**DOI:** 10.1101/2023.03.27.534486

**Authors:** Suman Saha, Priyanka Chakraborty, Amit Naskar, Dipanjan Roy, Arpan Banerjee

## Abstract

Ageing brain is associated with a slow drift in structural network properties over the lifespan accompanied by reorganization in neuromolecular interactions giving rise to changes in global functional markers. What are the guiding principles of the homeostatic mechanisms that maintain the desired performance of functional neural circuits and preserve brain health during healthy ageing? We hypothesize that an ageing brain alters two primary neurotransmitters, glutamate and *γ*-aminobutyric acid (GABA), responsible for excitation-inhibition regulation, concomitant with anatomical demyelination to preserve critical neural dynamics that are necessary to uphold optimal network performance. Thus, often observed re-organized functional connectivity with age by several investigations is a byproduct of this adaptive process. We demonstrate that the adaptive mechanism is driven by the tuning of glutamate and GABA concentration over a very slow time scale (lifespan) that can be estimated by tracking criticality from co-ordinated neural dynamics at a resting state via a biophysically driven computational framework, introduced as a computational microscope. We validate several empirical observations and model predictions across three independent aging cohorts using this computational microscope. One of the key mechanisms we discover is the reduction in local glutamate levels employed by brain regions to maintain a homeostatic balance with aging. This is further supported by the invariance of measures of global functional integration during the healthy ageing process.

## 1 Introduction

Ageing has a broad range of impacts on the healthy brain spanning multiples scales, ranging from microscopic level alteration in neurotransmitter levels [Cotman et al., 1987, Kaiser et al., 2005, Rojas et al., 2014, Puts et al., 2017, Maes et al., 2018, Roalf et al., 2020, Porges et al., 2021], degradation in long-range white matter tracts among brain regions [Samanez-Larkin et al., 2010, Coelho et al., 2021b] or gray matter volume (GMV) [Sullivan and Pfefferbaum, 2006], to the macroscopic level changes in functional coordination causing cognitive and behavioral impairments [Jolles, 1986, Benedetti et al., 1990, Li et al., 2020] and changes in functional integration mechanisms unfolding in brain networks [Hirsiger et al., 2016, Naik et al., 2017a, Luo et al., 2020]. Because of such multi-scale interactions, the effects of ageing on neural activity become complex, for example, some functions of a healthy ageing brain deteriorate largely, in contrast, others remain intact or even improve with age [Schaie, 1996, Naik et al., 2017a, Uddin, 2021, Aron et al., 2022]. So an overarching question arises, whether the age-related changes in neurobehavioral markers can be captured via an operational framework that ties up the myriad of complex interactions that spans multiple time-scales, e.g., neural field dynamics and neurotransmitter kinetics.

To conceptualize an operational framework for understanding age-related changes in the brain, one can delve into the structural and functional markers [Betzel et al., 2014, Petkoski et al., 2023] across the lifespan. Typically, the topology of a structural brain network can be quantified via measures from graph theory, such as modularity and global efficiency [Newman, 2006, Li et al., 2020, Coelho et al., 2021a]. Specifically, using a longitudinal design, Coelho and colleagues observed that modularity in a whole-brain structural network increases with age over a moderate-sized group of human participants (N=51). This general trend also survived for cross-sectional data from different publicly available cohorts of large sizes (N=174) used in this study. However, the connection probability computed across the entire brain didn’t change (Fig 1 a,b). An increasing number of studies have suggested that the functional connectivity during the resting state as a defining metric to capture the reorganization across lifespan ageing [Garrett et al., 2013, Betzel et al., 2014, Naik et al., 2017a, Thuwal et al., 2021, Pathak et al., 2022, Sastry et al., 2022]. Interestingly, the modularity remains preserved while the connection probability changes with age (Fig 1 c,d) when investigated over different data sets. To specifically address the dynamical characteristics of brain function, the mathematical measure of metastability is being increasingly used [Kelso, 2012, Deco and Kringelbach, 2016, Naik et al., 2017a]. Metastability can de ne the dynamic working point of a normal brain where it does not commit to any attractor state but remains perpetually prepared to internalize an inclement stimulus that drives to specific attractors [Pillai and Jirsa, 2017]. Thus, being in a metastable state allows the brain to maximize information processing [Haldeman and Beggs, 2005, Schneidman et al., 2006, Vázquez-Rodr guez et al., 2017]. Hence, any conceptual framework relating to the empirical observations of brain structure and function must be constrained to brain states that maximize metastability. Interestingly, we report that metastability is preserved across young and elderly (Fig 1); however, if the intermediate age range is taken into consideration, the relationship may be more U-shaped [Naik et al., 2017a, Naik et al., 2017b].

**Figure 1.**
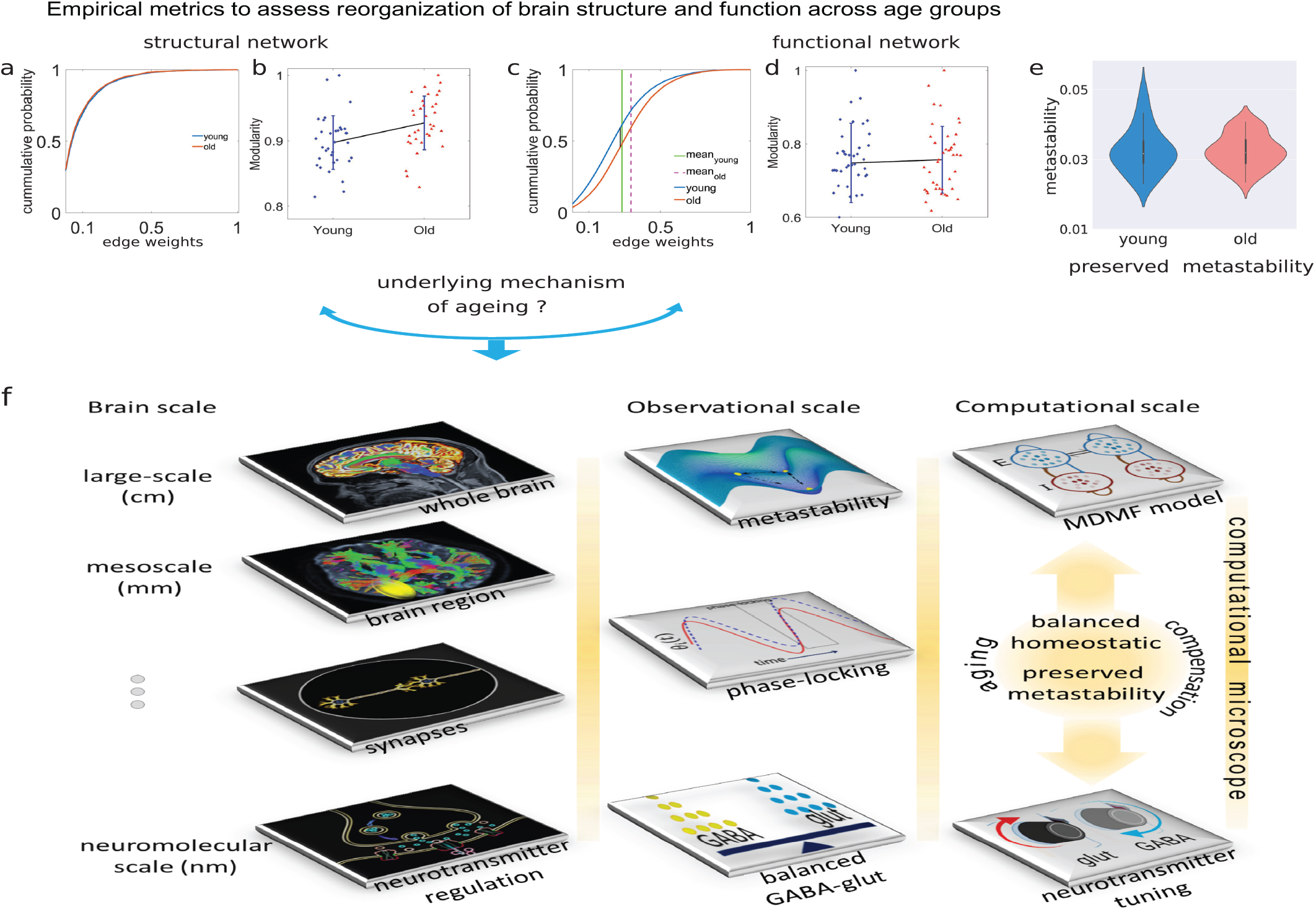
Observations from empirical data and various scales. Top row (a-e) shows age-dependent variance/invariance in the empirical data between two age groups. Observations on structural and functional networks are shown in (a,b) and (c,d). (e) A representative result for metastability derived from the empirical BOLD signals, showing no significant changes between the two age groups. (f) First column presents a schematic diagram of different brain scales. Individual tiles represent different brain scales. Second column shows observational scales, where metastability and phase locking are observed at the large-scale level of the brain, and glutamate-GABA levels are curated at the neuromolecular scale. Third column shows the computational microscope. The top tile shows that the MDMF model is placed on top of a real anatomical network, a large-scale computational model. The lower tile shows the optimal tuning of the two parameters at the neuromolecular level to maintain balanced homeostasis across whole brain.

In this study, we take the view that principles of neurocompensation underlying healthy ageing can be characterized by applying graph theoretic measures on structural and functional connectivity [Rubinov and Sporns, 2010]. Accordingly, we apply these measures to three cohorts of healthy ageing to identify the patterns of invariant functional segregation, integration, and resilience across healthy ageing. We also study the reorganization in the dynamic working point of the brain via the measure metastability [Deco et al., 2017], which can potentially characterize the change of homeostatic balance in brain dynamics across ageing trajectories. We hypothesized that metastable dynamics, observed in EEG/ MEG/ fMRI emerge from the multi-scale interactions involving neurotransmitter kinetics and excitatory-inhibitory membrane currents at the neural field level [Naskar et al., 2021]. Previous studies have demonstrated that regional E-I homeostasis determines the local excitability and critical firing rate ∼3*Hz* [Vogels et al., 2011, Deco et al., 2014, Vattikonda et al., 2016] of neural populations and controlled by brain’s intrinsic parameters, e.g., neurotransmitters. Hence, the combination of structural topology and controlling parameters such as excitatory-inhibitory (E-I) balance determine the critical brain dynamics [Marder and Goaillard, 2006]. Accordingly, we deploy a biophysically realistic computational model of whole-brain network which captures resting state activity as a function of synaptic coupling and neurotransmitter concentrations, Glutamate and GABA [Deco et al., 2014, Naskar et al., 2021], to understand the neuromolecular changes across lifespan ageing. Using metastability as the key constraining factor, we undertake a model inversion to predict the GABA, and Glutamate concentration changes over healthy ageing, which is weighed against empirical evidence obtained through magnetic resonance spectroscopy studies in the literature. Hence, the modeling framework can be considered as a computational microscope to reveal the neuromolecular-field level interactions of the nervous system, otherwise inaccessible using non-invasive neuroimaging technologies, and has the potential to track any neurodevelopmental disorders where the homeostatic balance of excitatory-inhibitory neurotransmitters is altered.

## 2 Materials and methods

### 2.1 Empirical data sources

Data sets for this study were obtained from 3 different sources. We acknowledge Berlin Centre for Advanced Imaging, Charité University Medicine, Berlin [Schirner et al., 2015] for data collection and sharing. We used the pre-processed data provided by the group. This dataset was collected by a 3T Siemens Tim Trio scanner with a 32-channel head coil (2mm isotropic voxel size). The pre-processed data from the Nathan Kline Institute (NKI) Rockland was obtained from the publicly accessible UCLA Multimodal Connectivity Database (UMCD) [Brown et al., 2012]. We acknowledge the Cambridge Centre for Ageing and Neuroscience (Cam-CAN), University of Cambridge, UK, for data collection and sharing. They used 3T Siemens Tim trio scanner with a 32-channel head coil (voxel size 3 × 3 × 4.4mm [Shafto et al., 2014]). We have pre-processed randomly selected particpants from similar age groups to Berlin and NKI data. Details of participants, data acquisition, and pre-processing are mentioned in the Supplementary Materials with a summary of demographics in Table 1.

**Table 1:**
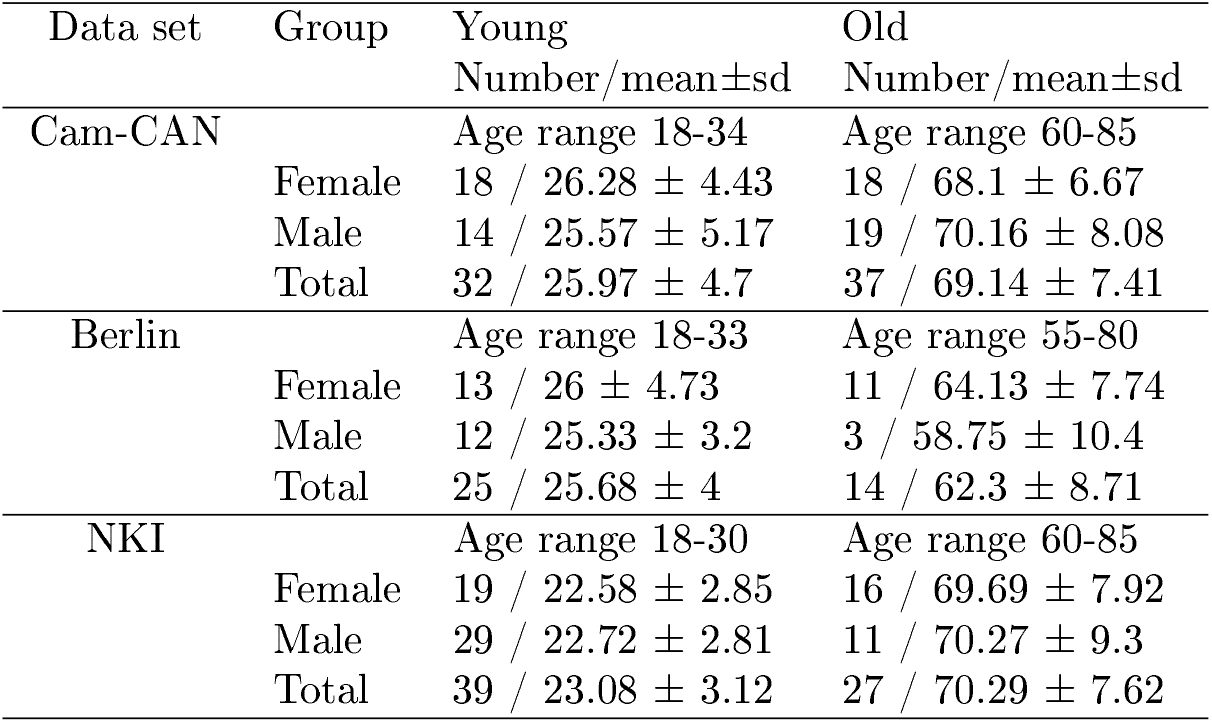
Demographic data across two different stages of adulthood in men and women.

### 2.1.1 Preparing SC and FC for network analysis

SC matrix is normalized by the maximum count of fibers, ranges 0 ≤ SC ≥ 1 for individual subjects obtained from diffusion weighted imaging (dWI). For FC (functional connectivity), we converted the FC matrix into a binary network by applying a threshold on the absolute values of node weights. Firstly, we normalized the absolute values of FC and then converted it into a binary matrix, where *FC*_*i,j*_=1, if |*FC*_*i,j*_| > *δ*, and *δ*=0.001 is a small threshold; otherwise, *FC*_*i,j*_=0. Next, we have used undirected binary network measures from the Brain Connectivity Toolbox (BCT) (http://www.brain-connectivity-toolbox.net) for our analysis. Weighted network measures are used for individual subjects with normalized SC matrices. We then tested for statistical significance in the changes of network properties across two age groups using independent t-test and Mann-Whitney U-test analyses for three data sets. Please refer to Fig. S1 in Supplementary Materials for details.

### 2.1.2 Network properties

The network properties were computed using BCT [Rubinov and Sporns, 2010]. The definition and description of the measures are:

1. Modularity [Newman, 2006], 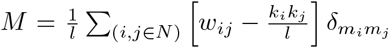, where l is the sum of weights in the network; *w*_*ij*_ is the connectivity weight between nodes *i* and *j*; *k*_*i*_, *k*_*j*_ are the weighted degrees of nodes *i* and *j*, respectively. Modularity gives network resilience and adaptability, measuring the degree of segregation. Communities are sub-groups of densely interconnected nodes sparsely connected with the rest of the network. In the case of a functional network, modularity signifies coherent clusters of functional modules.
2. Assortativity [Newman et al., 2006], 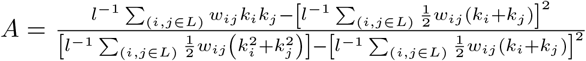, where *k*_*i*_, *k*_*j*_ are the weighted degrees of nodes *i* and *j*, respectively; *w*_*ij*_ is the connectivity weight between nodes *i* and *j*; *L* is the set of all edges within the network, and *l* is the total number of edges. Assortativity depicts the network’s resilience.
3. Global efficiency [Latora and Marchiori, 2001], 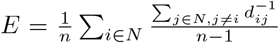, where *N* is the set of all nodes; *n* is the total number of nodes; *d*_*ij*_ is the weighted shortest path length between node *i* and *j*. It captures the integration property of a network.
4. Characteristic path length [Watts and Strogatz, 1998], 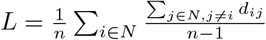, where *N* is the set of all nodes; *n* is the total nodes; *d*_*ij*_ is the weighted shortest path length between *i* and *j*. The characteristic path length for weighted graphs is an estimate of proximity. The global efficiency is the average of the inverse shortest path length. The global efficiency is computed on the node’s neighborhood and is related to the clustering coefficient.
5. Transitivity [Humphries and Gurney, 2008], or clustering coeffcient, is a measure of the tendency of the nodes to cluster together. High transitivity means the network contains communities or groups of nodes that are densely connected internally. Transitivity of a graph with degree sequence *k* is 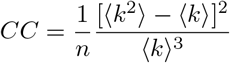, where ⟨*k*⟩ = 1/*n* Σ_*i*_ *k*_*i*_ is the mean degree and 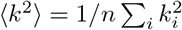 is the mean square degree.
6. Node betweenness centrality [Kintali, 2008]: The betweenness centrality of vertex *i* is: 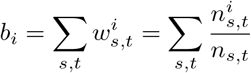 where by convention the ratio 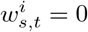 if *n*_*s,t*_ = 0. Each pair of vertex *s, t* contributes to the sum for *i* with weight 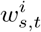 between 0 and 1, expressing the betweenness of *i* with respect to the pair *s, t*. We took (1/*n*) ∑_*i*∈*n*_ *b*_*i*_ to get the average node betweenness centrality. Betweenness centrality measures the extent to which a vertex lies on paths between other vertices. Vertices with high betweenness have considerable influence within a network under their control over information passing between others. If the vertices are removed, they will cost in disrupted communication, thus depicting the network’s resilience or vulnerability.

### 2.2 Descriptions of computational microscope

We use the multi-scale dynamic mean eld modelling framework (MDMF) [Naskar et al., 2021] to model the homeostatic balance of excitatory-inhibitory interactions that we hypothesized to be a key factor for the preservation and improvement of functional integration measured via topological and dynamic measures. The dynamics of a putative brain area is governed by a stochastic nonlinear dynamical system, i.e. the MDMF model which are connected via dWI derived connection topologies of each participant. Thus, the whole brain model of each area comprising an MDMF unit takes structural connectivity matrix as one of the inputs to generate simulated functional connectivity as an output that can be compared against empirical rs-FC to fit the neurotransmitter concentrations in an optimal way.

Two free parameters control the underlying dynamics of the MDMF-based whole-brain model, the concentration of GABA and Glutamate. Detailed expressions of how the concentration of GABA and glutamate can be obtained from population-level averaging from individual neuron-level synaptic concentrations are presented in our earlier study [Naskar et al., 2021]. Following similar lines of reasoning, we assume the whole brain-level modulations of GABA and glutamate are homogeneous at rest. Thus, the optimal levels of GABA and Glutamate can be estimated from a model inversion approach where the whole-brain dynamics is constrained according to the following conditions

(i) a normal healthy brain is at rest, exhibits criticality, and displays maximum metastability and (ii) the functional connectivity distance (FCD), a Euclidean distance between MDMF model generated-FC and empirical rs-FC is minimized. The concept is elaborated with detailed descriptions and gures; see Fig. S5 in Supplementary Materials.

#### 2.2.1 Multi-scale Dynamic Mean Field (MDMF) model

The whole-brain dynamics is described by the current-based MDMF model [Naskar et al., 2021]

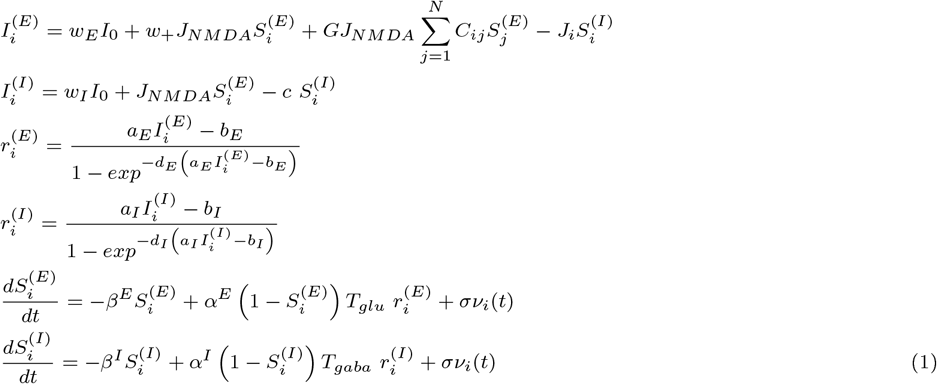

where, the subscripts *i, j* indicate brain areas, *i, j*=1,2, …, *N*. The total number of considered brain areas is *N*, where its value is different for different parcellation choices, e.g., *N* = 68 for Desikan-Killiany parcellation [Desikan et al., 2006], and 188 for Craddock 200 atlas [Craddock et al., 2012]. The superscripts *E* and *I* represent excitatory and inhibitory populations, respectively. 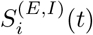 (*t*) denotes the average excitatory or inhibitory synaptic gating variables of the *i*^*th*^ region. The time-dependent gating variables are drawn by averaging the fraction of open channels of neurons. The two parameters, *T*_*glu*_ and *T*_*gaba*_, capture the neurophysiology of the excitatory and inhibitory neurotransmission process. Stochasticity is introduced by additive uncorrelated white Gaussian noise *ν*_*i*_ in two gating variables with intensity *σ* for individual regions. Further details of the model formulation and descriptions are available in the article by A. Naskar et al. [Naskar et al., 2021]. Default parameters are given in Supplementary Materials Table S2.

*J*_*i*_ denotes the local synaptic coupling strength from inhibitory to excitatory pools. The governing dynamics of the inhibitory feedback is,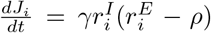. At the meaneld level, the biological complexity involved in the balance of E-I can be captured grossly using the mathematical implementation of the inhibitory plasticity rule [Hellyer et al., 2016]. An inhibitory plasticity rule represents changes in *J*_*i*_(*t*) (synaptic weight) to ensure that the inhibitory current clamps to an excitatory population. Thus, homeostasis is achieved by the dynamics of local inhibitory weights *J*_*i*_(*t*) as a function of time, such that the firing rate of the excitatory population is maintained at the target firing rate *ρ* = 3*Hz*. The chosen target firing rate emerges when E-I balance is achieved. *γ* is the learning rate in *sec*.

#### 2.2.2 FC distance (FCD) measure

Model-based synthetic neural activity was converted into BOLD time series by Balloon-Windkessel hemodynamic algorithm [Friston et al., 2003]. We generate a model FC for each subject from the model-based BOLD signals deriving a Pearson correlation between each pair of brain regions. To compare the model FC with empirical FC, we calculated the FC distance (FCD)for each combination of GABA and glutamate. FCD for a total of *n* regions is calculated using,

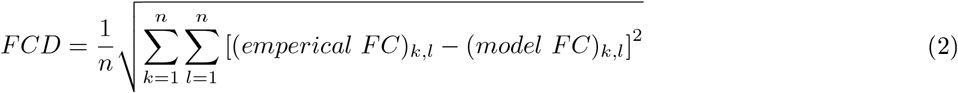

#### 2.2.3 Global coherence and metastability

Recent computational brain network models [Deco et al., 2017, Surampudi et al., 2019] have predicted (for instance) that the human brain operates at maximum metastability during the resting state. It is manifested as an optimal information processing capability and switching behavior [Werner, 2007, Deco et al., 2009, Shanahan, 2010]. Also, the existing cognitive ageing theories are explained better with the concept of metastability [Naik et al., 2017a] and used to explore changes in the physiological substrate over ageing [Naik et al., 2017a, Naik et al., 2017b]. We have chosen metastability as one of the optimal conditions to connect the parametric role to the functional organization.

First, an instantaneous order parameter *r*(*t*) is calculated.

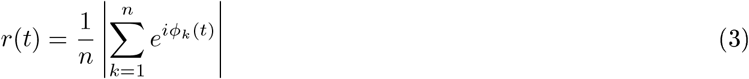

where, *ϕ*_*k*_(*t*) is the instantaneous phase of *k*^*th*^ brain region. The value of *r*(*t*) lies between 0 ≤ *r*(*t*) ≤ 1; *r*(*t*) = 1 representing complete synchronous, *r*(*t*) = 0 for desynchrony, and (0 *< r*(*t*) *<* 1) representing partial synchrony among all brain regions. Hence, *r*(*t*) indexes the Global coherence among a system of oscillating brain regions.

We can measure the temporal variability of *r*(*t*) to capture the fluctuations in global spatial synchrony within the whole brain over time. Variation in *r*(*t*) indicates the fluctuation among collective states (such as coherent or incoherent) over time, which is indexed by metastability. Metastability [Naskar et al., 2021] is used to capture the temporal fluctuation, estimating the tendency of a brain region to deviate from the coherent manifold. Thus, it could be a better measurable unit for observing spatial cohesion for a short period (maybe a few hours) and may also serve as a key constituent for observing a gradual change in lifespan brain neural dynamics on a parametric plane (e.g., GABA, glutamate plane) over a very slow time scale, such as the lifespan. Metastability (**M**) can be calculated by taking the standard deviation of r(t) over time: 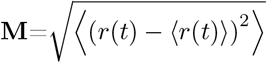, where ⟨.⟩ denotes time average.

#### 2.2.4 Estimation of optimal GGC and GGR

We use two measures, functional connectivity distance (FCD) and metastability (M), to estimate the optimal values of glutamate (*T*_*glu*_) and GABA (*T*_*gaba*_) concentrations in units of mMol. For each value of the two parameters, we compute *FCD*(*T*_*glu*_, *T*_*gaba*_) and **M**(*T*_*glu*_, *T*_*gaba*_) to generate a two-parameter space. We determine the optimal conditions as the (i) minimum of *FCD*(*T*_*glu*_, *T*_*gaba*_), and (ii) maximum of **M**(*T*_*glu*_, *T*_*gaba*_). We first estimate 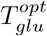 for different values of *T*_*gaba*_ that satisfy both conditions. Then, we estimate 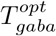among the obtained results by imposing similar conditions. This way, we obtain each pair of optimal concentrations, *GGC* = (*T*_*glu*_, *T*_*gaba*_)^*opt*^ in mMol. We calculate the GABA-glutamate ratio (GGR) by taking their ratio as *GGR* = (*T*_*glu*_*/T*_*gaba*_)^*opt*^ for individual subjects. Finally, we perform a two-sample t-test to determine if there are any changes in the estimated *GGC* and *GGR* between the two age groups in the three datasets.

## 3 Results

We present the results from this study in two parts. First, we report the observations from network analysis performed on structural and functional imaging data depicting how alterations of graph theoretical metrics represent neurocompensatory mechanisms. The empirical observations were validated in 3 independent data sets. We purposefully chose to analyze two age groups in the boundaries of young (18-34) and elderly (60-85) to characterize the salient changes in network properties across the lifespan, rather than a lifespan continuum followed in our earlier studies on lifespan ageing [Sahoo et al., 2020]. Second, we apply the computational microscope to uncover the neuromolecular interactions that preserve homeostasis across lifespan.

### 3.1 Reorganization of network architecture and dynamics across lifespan

We have measured network properties that capture segregation, integration, and resilience of structural and functional networks to understand the age-dependent change in the brain networks [Rubinov and Sporns, 2010]. The network metrics were computed using Brain Connectivity Toolbox (http://www.brain-connectivity-toolbox.net), from structural connectivity (SC) and resting-state functional connectivity (rsFC), presented in upper panels, Figs. 2(a-f) and middle panels, Figs. 2(g-l), respectively. Statistical significance in the metrics between two age groups was determined by independent t-test (for Cam-CAN and NKI datasets) and Mann-Whitney U-test (for Berlin dataset). The structural and functional network changes are observed based on segregation (Figs. 2(a,b) and 2(g,h)), integration (Figs. 2(c,d) and 2(i,j)) and resilience (Figs. 2(e,f) and 2(k,l)) of the networks. Figures 2(m,n) present violin plot for metastability, measured for individuals from the two groups. To assess the changes in neural coordination dynamics among brain areas across lifespan we applied the measure of metastability.

**Figure 2.**
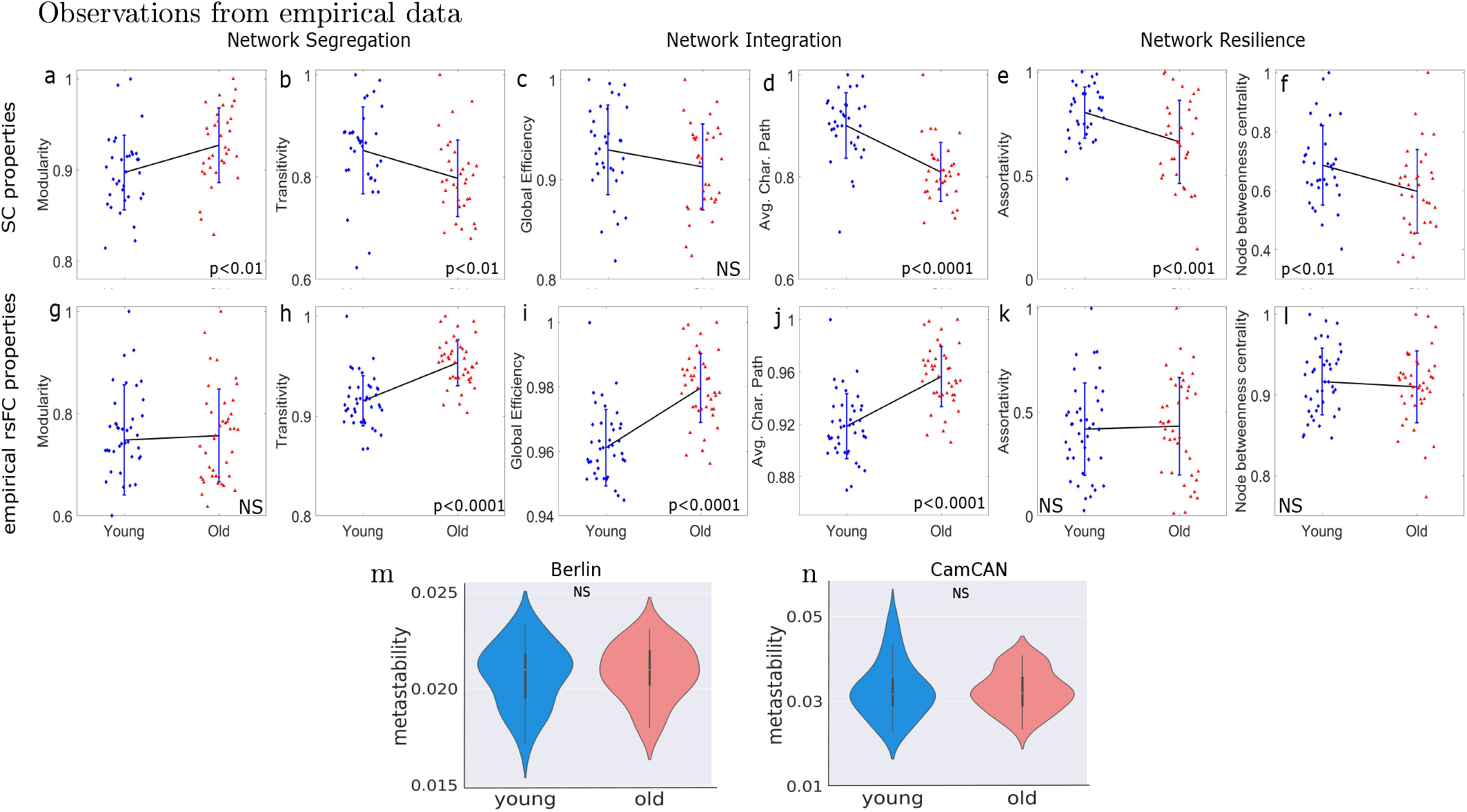
Changes in SC and functional network properties in Cam-CAN data set. Top panels (a-f) present the anatomical network properties of individual subjects from two age groups. (a) Structural modularity is increased, and (b) transitivity shows a decreasing trend. It can be interpreted as declined structural segregation. Due to loss in white-matter tracts in the ageing brain, the connections across modules become more sparse. Thus, segregated SC resulted in a decreased inter-regional exchange of excitation. Though (c) global efficiency remains invariant, (d) significantly decreased average characteristic path length indicates a decline in network integration, which has a direct impact on the network resilience, conformed by (e) assortativity, and (f) node betweenness centrality comparing the two age groups. Next, Functional network properties are shown in the second row (g-l). (g) Functional modularity remains unaltered with age, implying an average number of functional modules remains intact. However, specific brain regions could vary within a module in the two age groups. A significant increase (*p <* 0.001) is observed in (i) global efficiency and (j) average characteristic path length. Increased global efficiency and average characteristic path length in the elderly group signify an increased functional network integration. No significant changes are seen in (k) assortativity and (l) node betweenness centrality between the two groups, which describe unaltered functional resilience in ageing brain. To visualize alteration patterns between the age groups, we join the two means by a black line, where the standard deviations from the mean are shown in blue lines. The independent t-test analysis determines significant changes (no changes). Network properties are derived using Brain Connectivity Toolbox [Rubinov and Sporns, 2010]. Lower panels (m,n) show metastability derived from the empirical BOLD signals of Berlin and Cam-CAN data sets. No statistically significant difference is found in the metastability between the two age groups in both data sets.

#### 3.1.1 Segregation: Modularity and Transitivity

Segregation is a network phenomenon where dense connectivity is observed within the sub-modules of a large network; however, sparse connections among the sub-modules exist [Rubinov and Sporns, 2010, Newman, 2006]. To characterize network segregation, we computed modularity and transitivity of structural and functional networks from structural connectivity and rs-FC matrices, shown in Figs. 2(a,b) and 2(g,h), respectively. Modularity signifies how the network is densely connected among the areas physically within a sub-cluster but has sparse interactions across the segregated sub-clusters. A network’s transitivity or clustering coefficient measures how nodes cluster together. Lower transitivity means the network contains sparsely connected dominant sub-modules. Thus, both measures qualify as network segregation measures.

Increased modularity [Li et al., 2020] [shown in Fig. 2(a)] and decreased transitivity [in Fig. 2(b)] along lifespan was observed while applying these measures on structural connectivity matrices. Earlier studies have shown segregated structure is an outcome of the factors affected mainly by age, e.g., white-matter fiber tracts reductions [Coelho et al., 2021b, Kristanto et al., 2020, Davis et al., 2009, Petkoski et al., 2023] and degradation in long-range interareal connections [Betzel and Bassett, 2018], so this is a reconfirmation of previous results across 3 data sets.

Next, modularity and transitivity were applied in whole brain rs-FC matrices to characterize functional segregation. Invariance in modularity [Fig. 2(g)] computed from rs-FC implies no change in the number of functional modules or clusters evolved from the resting state BOLD activity in the large-scale brain network. It is observed that the transitivity of structural network was decreased [Fig. 2(b)] in elder subjects, whereas transitivity of functional network increased with age [Fig. 2(h)]. Significantly changed transitivity of FCs in elderly subjects indicates rewiring in the functional connections while preserving modularity, i.e., keeping functional modules intact. An ageing brain might calibrate controlling parameters to maintain a self-sustaining pattern in brain dynamics preserving functional modules to avoid functional decline due to altered structure. However, the regions within individual functional clusters or modules may di er across subjects and age groups over time.

#### 3.1.2 Integration: Global efficiency

Structural integration signifies that paths are sequences of distinct areas and links. It represents potential routes of information flow between pairs of brain areas [Rubinov and Sporns, 2010]. Network integration is observed using global efficiency and characteristic path length from SCs and FCs, shown in Figs. 2(c,d) and 2(i,j), respectively. Structural integration [Rubinov and Sporns, 2010] declines with age as suggested by a decreasing trend in global efficiency [Li et al., 2020] and significantly decreased characteristic path length [Figs. 2(c,d)] in elderly subjects. In general, the characteristic path length is primarily influenced by long paths (infinitely long paths are an illustrative extreme), while the short paths primarily influence global efficiency [Rubinov and Sporns, 2010]. On more extensive and sparer networks, paths between disconnected nodes are defined to have infinite length, and correspondingly zero efficiency, thus affecting long- and short-range information flows across the whole brain.

Functional integration in the brain can be interpreted as the ability to combine specialized information from distributed brain areas rapidly [Rubinov and Sporns, 2010]. Paths in functional networks represent sequences of statistical associations and may not correspond to information flow on anatomical connections [Rubinov and Sporns, 2010]. Precisely, a functional network’s global efficiency measures a network’s ability to transmit information at the global level [Wang et al., 2010]. Increased functional integration, as observed from an increasing trend in global efficiency and average characteristic path length in older groups [see Figs. 2(i,j)], might be associated with the compensatory process against the decline in structural integration [Figs. 2(c,d)], which enhances the communicability or the ability to combine technical information from distributed cortical circuits. The relationship between structure and functions may not necessarily be one-to-one linear mapping, as intrinsic biological parameters largely influence the emergence of collective cortical dynamics.

#### 3.1.3 Resilience: Assortativity and betweenness centrality

Resilience refers to the ability of brain networks to recover and preserve functions under inclement structural degradation or insults [Rubinov and Sporns, 2010]. Some studies have shown that certain brain areas are more prone to overall disruption of function, but several nodes remain unaffected by random perturbations [Achard et al., 2006]. The fewer the number of such influential nodes, the more resilient is the network. The commonly used metrics of network resilience are assortativity, and node betweenness centrality [Rubinov and Sporns, 2010]. Networks with a positive assortativity coefficient likely have a resilient core of mutually interconnected hubs. In contrast, a negative assortativity coefficient is likely to have widely distributed and consequently vulnerable hubs [Rubinov and Sporns, 2010].

Assortativity and node betweenness centrality of SCs and FCs are shown in Figs. 2(e,f) and 2(k,l), respectively. Resilience of structural network [Figs. 2(e,f)] is significantly decreased in the older group, as observed from a significant decline in assortativity and betweenness centrality in Figs. 2(e) and 2(f), respectively. The assortativity coefficient is a correlation coefficient between the strengths (weighted degrees) of all nodes on two opposite ends of a link. Decreased assortativity indicates a reduced tendency to link nodes with similar strengths. It reconfirmed our previous observations on reduced integration and increased segregation in structure, which could occur due to the loss of connectivity strengths (network became sparse, weighted degrees of individual nodes were decreased) and deterioration in long-range white-matter tracts connecting distant brain regions [Petkoski et al., 2023].

On the other hand, resilience of the functional network from rs-FC, measured at the level of individual subjects using assortativity (Fig. 2(k)) and node betweenness centrality (Fig. 2(l)), remains unaltered, essentially no significant changes (*p*=0.6 and *p*=0.45), between two groups. It can be interpreted as a functional compensation among nodes of ageing brain against the deteriorated structural resilience (Figs. 2(e,f)), thus upholding functional resilience.

We did test-retest validation on two other data sets, Berlin and NKI data, and the results are included in Fig. S1 in Supplementary Materials. We observed quite a similar pattern of alteration and re-organization in empirical SC and rsFC properties, as seen in Cam-CAN data. Though no statistical significance is seen in the network properties for the Berlin data set (which could be due to comparatively fewer subjects in the two groups), but it shows similar alteration patterns as seen in the other two data sets.

#### 3.1.4 Dynamic working point: Metastability

Metastability indexes the perpetual state of transition observed in brain signals without committing to a speci c attractor state [Bressler and Kelso, 2001, Tognoli and Kelso, 2009, Tognoli and Kelso, 2014] and has been used as a metric to capture the dynamic working point of the brain at rest by several researchers [Deco and Kringelbach, 2016, Deco et al., 2017, Naskar et al., 2021]. There are several advantages of using metastability such as, the metric can be used to quantify the tendencies of neural systems to operate at segregative and integrative modes [Tognoli and Kelso, 2014], characterize the state of criticality where any dynamical system remains on the verge of phase transition thus enhancing information processing capabilities [Deco et al., 2017, Cocchi et al., 2017]. Finally, the phenomena can be studied via generative models of neural dynamics, and hence allows the researchers to gain mechanistic insights into network mechanisms generating brain signals [Breakspear, 2017, Roberts et al., 2019, Naskar et al., 2021].

Metastability is defined as the standard deviation of the order parameter, the vector sum of the individual phase of network nodes typically observable in empirical BOLD time series obtained through Hilbert transforms. Violin plots of metastability of young and elderly subjects are shown in Fig. 2(m) for Berlin data and Fig. 2(n) for Cam-CAN data. First, we performed a z-transform of BOLD signals of individual brain areas for each subject. Next, the instantaneous phase of each region was determined using the Matlab function ‘hilbert.m’. These two steps are repeated for individual subjects. We checked statistical variations between the two age groups using an independent t-test for Cam-CAN data and Mann-Whitney U-test for Berlin data. We observed no significant changes in the metastability between the two age groups, suggesting that despite structure changes, the dynamic working point remains unaltered in elderly subjects.

#### 3.1.5 Summary of observations on empirical data

We employed network analysis tools to structural and functional connectivity matrices in young and aged cohorts from 3 different data sets and report that although there are overarching changes in network structure, the measures of segregative and integrative brain function are preserved across ageing. This led us to hypothesize that a mechanistic approach taken by the ageing brain to achieve function preservation may be via changing the neurotransmitter concentrations to maintain a homeostatic equilibrium between excitatory and inhibitory current dynamics. Previous studies have proposed that a homeostatic balance can be maintained via maximizing metastability [Naskar et al., 2021] - which did not show any change between young and the elderly groups (Fig 2). Thus, the goal of the remaining results section is to identify the age-associated changes in GABA-Glutamate concentrations while keeping the dynamic working point (i.e., metastability) preserved.

### 3.2 Estimated glutamate-GABA concentrations from the MDMF model inversion

Estimated glutamate-GABA concentrations (GGC) and GABA/glutamate ratio (GGR) at the level of individual subjects from the two age groups are shown in Fig. 3, results from Berlin data set in Figs. 3(a-e), Cam-CAN data in Figs. 3(f-h) and NKI data in Figs. 3(i-k).

**Figure 3.**
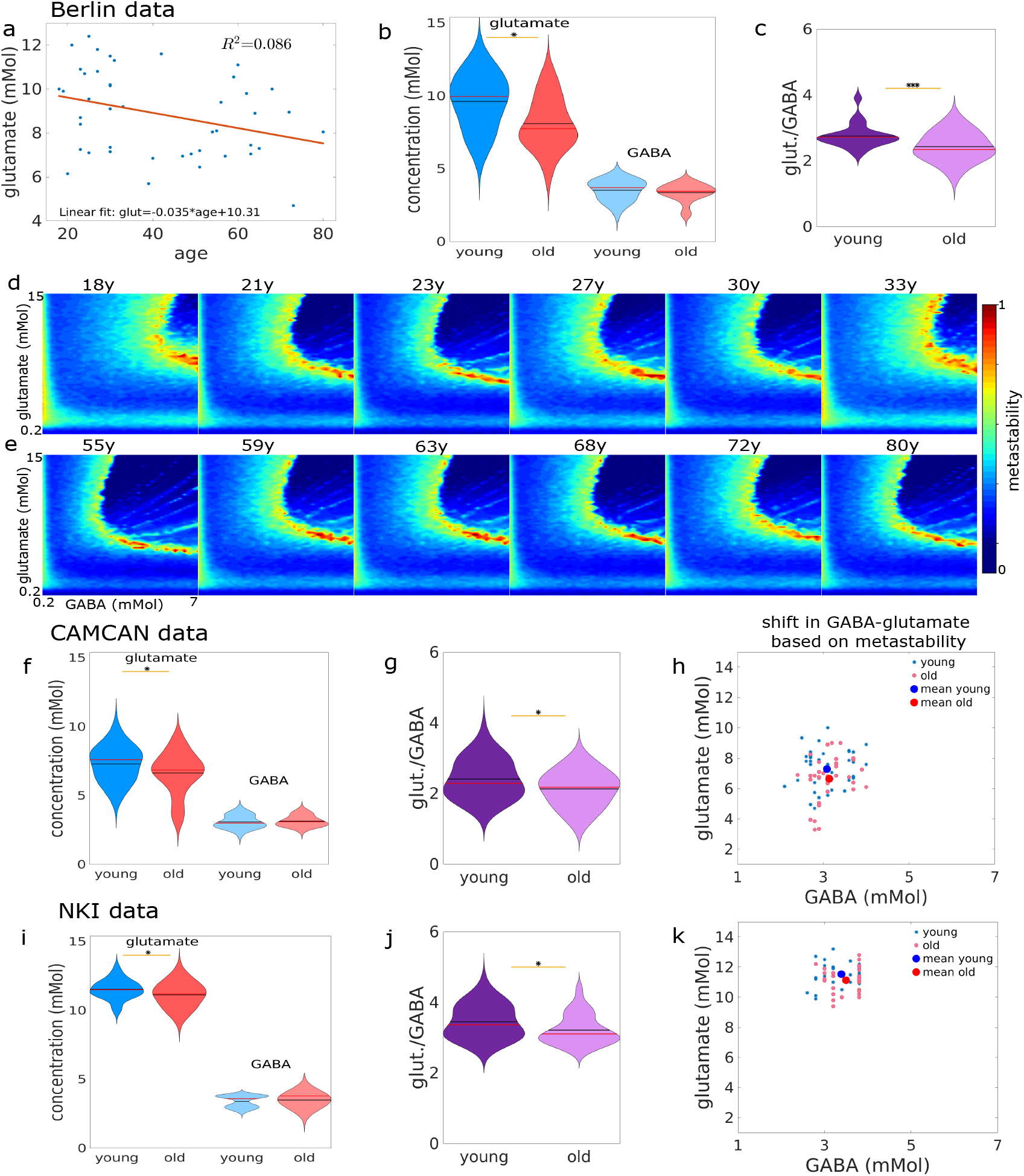
Subject-wise alteration in GGC and GGR over age. MDMF model inversion was used to estimate GABA-Glutamate concentration (GGC) and GABA-Glutamate ratio (GGR) in 3 cohorts on a participant-by-participant basis (a) Linear fitting (*R*^2^=0.086) shows a negative correlation between glutamate and age. Glutamate decreases across age by a decay rate = 0.035. GGC and GGR in the two age groups are shown in (b) and (c). The ageing significantly changes glutamate (*p*=0.042), but GABA remains almost invariant. A significant decay in GGR is observed (d,e). The metastability of individual subjects is shown for the Berlin data set. The optimal values of glutamate and GABA are chosen where the model predicts empirical FC with better accuracy, i.e., with minimum FC distance. In addition, the collective brain dynamics operate at maximum metastable state. A shift in the maximum metastability (indicated by darker red) on the two-parameter space is evidenced during the healthy ageing process. Next, Cam-CAN data was analyzed and shown in (f-h). (f) GGC is shown using violin plots for two age groups. A significant (*p*=0.04) decrease in glutamate is observed in elderly subjects, and GABA levels remain invariant in the two groups. (g) A decline (*p*=0.028) in GGR can be evidenced, similar to the result obtained in Berlin data. (h) Shift in the levels of the metabolites are shown on the two-parameter plane based on optimal conditions, i.e., maximum metastability and minimum FCD. The shift in two neurotransmitters is the mechanism of the ageing brain to adapt to age-related changes. The adaptive mechanism provides a robust and general behavior to maintain neural activity intact and sustain the critical ring rate (excitatory, ∼ 4Hz). Similar tests are performed on the NKI-Rockland data set, where we plot GGC in (i), GGR in (j), and optimal values of all subjects on the GABA-glutamate parametric plane in (k). * for *p* < 0.05, and ** for *p* < 0.002.

Figure 3(a) depicts the changes in glutamate levels across ages. A significant declining trend is evidenced in the glutamate concentration with aging. A linear regression fitting is used to find a linear relationship between glutamate and age (*R*^2^ = 0.086). The linear fit shows that glutamate negatively correlates with increased age (slope= − 0.035). Figure 3(b) shows violin plots of estimated concentrations of the two metabolites in two age groups, where glutamate level is decreased significantly (*p <* 0.05), but GABA level remains invariant. Though, a prominent decrease in GABA-glutamate ratio is observed [Fig. 3(c)] from the model-estimated results between two age groups with a statistical significance of *p* = 0.0015. Figures 3(d,e) show few representative subject-wise changes in parameter space - metastability and its relation with *T*_*glu*_ and *T*_*gaba*_, where the color bar represents normalized metastability value. Interindividual variability is manifested as islands of metastability maximization in the two-parameter space of *T*_*glu*_ and *T*_*gaba*_ in representative subjects in chronological order (younger to older, 18 to 80 years).

Further, we have performed a similar analysis on the Cam-CAN data set, presented in Figs. 3(f-h). Figure 3(f) shows glutamate and GABA concentrations in young and old groups. Glutamate level is decreased in older groups, where GABA remains almost unchanged. A notable change in the ratio is observed (*p <* 0.03) in Fig. 3(g). To show the shift in glutamate and GABA levels in elderly subjects, we have plotted the estimated optimal concentrations of each subject in the two-parameter plane in Fig. 3(h). The average values of glutamate and GABA for the younger and older group are shown in the black star and green circle, respectively. The scattered blue stars and red circles are the estimated glutamate and GABA values for individual subjects, separating into young and older groups. The three data sets found a similar pattern of altered GABA glutamate levels with age.

We have performed a similar analysis on the NKI data set, presented in Figs. 3(i-k). Figure 3(i) shows glutamate and GABA concentrations in young and old groups. Glutamate level is decreased in older groups, where GABA remains almost unchanged. A notable change in the ratio is observed (*p <* 0.03) in Fig. 3(j).To show the shift in glutamate and GABA levels in elderly subjects, we have plotted the estimated optimal concentrations of each subject in the two-parameter plane in Fig. 3(k).

The altered level of glutamate concentrations and glutamate-GABA ratio can be interpreted from the neural dynamics viewpoint, correlating cortical cohesion (e.g., preserved metastability) with age-related factors, such as increased segregation and decreased integration and resilience in structure, functional network with decreased segregation, increased integration and invariance in resilience. An ageing brain shifts the optimal operating point by reducing glutamate level to compensate for age-related variability in the collective dynamics and avoids functional degradation. We observed that re-organizing FCs results from the underlying brain dynamics when the two neurotransmitters control the intrinsic compensatory processes to maintain an equilibrium in the E-I ratio at rest, where the excitatory firing rate sustains at a critical range of 2.55-3.65Hz. The whole brain cohesion pattern remains intact, i.e., preserved metastability in healthy ageing.

## 4 Discussion

Several critical reviews have highlighted the urgent need for tools that can leapfrog studies of neuroscience from a scale-specific biological observation to explain ongoing behavior towards the development of comprehensible multi-scale models that can generate an understanding of the interactions among multiple organizational scales [Rosen, 1991, Frackowiak and Markram, 2015, D’Angelo and Jirsa, 2022]. This study focuses on a whole-brain computational model that captures the interactions between two observational levels of neural complexity, ultra-slow BOLD-fMRI dynamics, and neurotransmitter kinetics. The precise interactions transcending the biological scales of observations allow the estimation of critical markers such as Glutamate-GABA concentrations from model inversion techniques across healthy human ageing. First, we illustrate that functional segregation, integration, and resilience measures can be preserved and sometimes enhanced across the healthy ageing process, even though similar metrics applied on structural network topology point out a gross degradation. Second, we identify a key invariant that captures the dynamic working point of the brain, as well as provides a link to model the interaction between neurotransmitter kinetics and emergent metastable dynamics of the local field with aging. Finally, we could predict the optimal concentration changes in GABA and glutamate associated with healthy aging by model inversion under the constraint of an optimized dynamic working point. Thus, our study develops a computational microscope based on a biophysically realistic and sufficiently detailed generative neural mass model. The advantage of using this model is two-fold, reconciling observational accounts of metabolite mapping with mechanistic insights of neuronal network workings and possibly setting up future studies that can use this framework to predict the onset of neurodevelopmental and neuroinflammatory disorders where the local E/I balance is crucially perturbed. We discuss these issues in more detail in the remaining sections.

### Functional neurocompensation along ageing trajectories

The most salient feature of our empirical network analysis is that over the lifespan, a maintenance and sometimes improvement were observed in measures of segregation, integration, and resilience applied on resting state functional connectivity (rs-FC), although the same metrics computed from structural connectivity (SC) deteriorates with age (Fig 2). Our observations are in line with earlier reports on age-related structural changes [Coelho et al., 2021a, Luo et al., 2020] and functional re-organization [Song et al., 2014, Zhang et al., 2021, Meunier et al., 2009]. In the field of ageing research, such phenomena come under the idea of neurocompensation [Reuter-Lorenz and Cappell, 2008, Park and Reuter-Lorenz, 2009] by which the brain areas improve upon their segregative and cooperative behavior under inclement structural decline [Daselaar et al., 2015, Pathak et al., 2022, Petkoski et al., 2023]. Since increased metastability means the system is essentially situated in a transient state where it can have more opportunities of choosing an attractor, this can also be interpreted as a state where information processing capabilities increase, leading to higher flexibility [Naik et al., 2017a]. Interestingly, metastability calculated from empirical BOLD signals was unaltered between the young and aged groups. This can be interpreted as the preservation of information processing across young and elderly cohorts, who, for the most part, are capable of complex cognitive functions (although showing a decrease in performance indices over age, see [Thuwal et al., 2021]) emerge from underlying compensatory mechanisms that preserves the global dynamical complexity of the brain. From the perspective of the current study, we use this observation as a constraining tool for model inversion using the MDMF model [Naskar et al., 2021] to estimate the Glutamate and GABA concentration changes across ageing trajectories.

### Insight from computational microscope: ageing associated decrease in Glutamate levels

A key contribution of the present study was the development of a framework that could estimate the local synaptic GABA, Glutamate concentrations (GGC) and GABA, Glutamate ratio (GGR) from non-invasive fMRI. We could predict that age negatively correlated with glutamatergic regulation, which was reduced with healthy ageing, in concurrence with earlier evidence [Davis and Himwich, 1975, Benedetti et al., 1990, Suri et al., 2017, Hu et al., 2013]. The alteration pattern in glutamate was consistent with previous age-related experiments [Suri et al., 2017, Hu et al., 2013], where glutamate was measured to be lower in older participants. A study [Liguz-Lecznar et al., 2015] concluded from similar observation that ageing down-regulated glutamate content and decreased the expression of several presynaptic markers involved in E-I balance. Increased excitability in older individuals was speculated to underlay the increased prevalence of certain neurological conditions with age, such as epilepsy [Staffaroni et al., 2018]. A study reported a similar negative correlation with glutamate contents in the ageing brain in the motor cortex [Kaiser et al., 2005], but a positive correlation concerning other metabolites, such as glutamine, N-acetyl aspartate, and creatinine. They further related the reduction of glutamate concentrations in the motor cortex with neuronal loss/shrinkage with ageing [Kaiser et al., 2005]. Studies in rodent brains also reported a similar reduction in glutamate associated with the healthy ageing process [Davis and Himwich, 1975, Benedetti et al., 1990]. Nevertheless, a review [Segovia et al., 2001] on the critical perspective of ageing and development on glutamatergic transmission had categorically shown the effects of ageing based on the shreds of evidence from various in-vitro and in-vivo studies on rodent brains, which led to contrasting conclusions. Few studies [Fornieles et al., 1986, Donzanti and Ung, 1990] had reported no significant changes in glutamate content in the dorsal prefrontal cortex, sulcal prefrontal cortex, temporal cortex, and medial prefrontal cortex during ageing. On the contrary, studies [Wallace and Dawson, 1990, Saransaari and Oja, 1995] had also reported significant changes at 20 50% in the glutamate contents in the frontal cortex, hippocampus, and cerebral cortex with ageing. Thus, for future work, it will be advisable to have a prediction of the more heterogenous distribution of metabolites in the brain which we discuss further in the following sub-section.

One crucial understanding that emerges from this work is that ageing affected GABA differently than glutamate, even though we varied GABA-glutamate concentrations homogeneously across brain areas. The estimated glutamate content was significantly reduced, whereas GABA showed no such variability with ageing. A similar pattern of altered metabolites was observed earlier in the context of ageing [Hu et al., 2013]. In contrast to our observation, studies [Rojas et al., 2014, Puts et al., 2017] have detected age-dependent reduction in GABA by magnetic resonance spectroscopy (MRS) studies in human subjects in the motor, visual, auditory, somatosensory areas, and the perisylvian region of the left hemisphere, leading to an abnormal information processing. It is unlikely that MRS gives a better accurate reflection of glutamatergic activity than GABAergic activity [Stagg et al., 2011] since GABA MRS acquisitions do not allow us to separate resonances from glutamate and glutamine. Most studies reported them as a composite measure of Glx. Finally, much fine-tuning of glutamatergic activity occurs via modulation of the NMDA receptors, which are invisible to MRS. MRS can detect a neurochemical concentration within a localized region (typically in the order of 2 × 2 × 2 cm^3^) of tissue [Stagg et al., 2011]. Studies suggest that MRS can not distinguish between separate functional pools of GABA as they might be more tightly bound to macromolecules than others, rendering them less “visible” to MRS [Floyer-Lea et al., 2006, Stagg et al., 2009]. In that case, our framework could be a supporting tool to look into the age-related fluctuations in the metabolites at rest and task-evoked activity, besides MRS-driven data, within a localized area and the whole-brain level.

A study [Gonen et al., 2020] using MRS on glutamate and GABA of the PCC/precuneus documented that glutamate and GABA did not significantly correlate with age on their own, but their ratio did. We found similar significant decay in GGR in elderly subjects. Though the declining pattern was more prominent in GGR than GGC, there was still significant decay in glutamate level in the three datasets. Ageing brain also can be viewed as a shift in the homeostatic state between local E-I balance mechanisms, which pitches GGR as a relevant metric to track rather than individual GGCs. A previous study [Liguz-Lecznar et al., 2015] showed how Glutamate-GABA equilibrium was altered in aged mice cortex, influencing cortical plasticity. We conclude that the E-I balance remains unaffected due to compensations in the ratio of the two neurotransmitters. Thus, the interplay between the tuned metabolite levels and cortical plasticity provokes a gradual functional re-organization in the ageing brain over a prolonged time (lifespan), preserving measures such as metastability.

### Limitations and future directions

While the computational microscope we have argued in this article gives a bird’s eye view of the neuromolecular mechanisms at the whole-brain level. Nonetheless, it is still more complex since it fails to capture spatial heterogeneity of metabolites in the cortex. Some potential limitations to our findings are as follows: (i) An earlier observation [Rozycka et al., 2019] on a rat brain experiment showed alteration in GABAergic and glutamatergic behavior effected by age was highly probable to occur in a presynaptic mechanism than in a postsynaptic mechanism. On the other hand, our method fails to curate data at a level of pre- or postsynaptic mechanisms of GABA-glutamate regulation [Pitler and Alger, 1994, Rozycka et al., 2019]. (ii) Uptake and release of glutamate [Saransaari and Oja, 1995] can not be separated using the present method. The model does not explicitly tell about the two major sub-types of GABA, i.e., GABA_A_ and GABA_B_. (iii) Measure of total GABA concentration remains unclear whether the GABA signal represents cytoplasmic, vesicular, or free extracellular GABA [Stagg et al., 2011].

The first limitation of heterogeneity of GABA-Glutamate not being captured can be addressed in further developments of the proposed hypothesis. An atlas of whole brain metabolite map from future MRS studies can be used to tune the MDMF model for better predictions of resting state BOLD dynamics. This would also require the implementation of a more advanced model inversion technique beyond the 2-parameter (GABA, Glutamate concentrations for this manuscript) fitting with optimized metastability and FCD used here. In other words, machine learning models such as Bayesian model inversion can replace this step. Future research can also explore how the imbalance in E-I homeostasis can occur due to disrupted neurotransmission that shapes neurophysiology differently for healthy brain and distinctively in various neural and mental disorders. The given framework can be a potential tool, besides MRS, in studying patient data against healthy controls keeping GGC as a primary constituent in the development of neuropathological conditions [Levy and Degnan, 2013], such as neural migration disorder [Luhmann et al., 2015], Parkinson [O’Gorman Tuura et al., 2018], Alzheimer [Huang et al., 2017], attention deficit hyperactivity disorder [Edden et al., 2012] and autism [Rojas et al., 2015, Purkayastha et al., 2015]. Earlier research documented how the alteration in the metabolites was helpful in early diagnosis of AD [Huang et al., 2017] or targeted increase in GABA release increase inhibition and effectively treat depression [Myers et al., 2014], by balancing E-I ratio. Thus, identifying neuromolecular interactions with synaptic dynamics can fill the gap towards developing more sophisticated therapeutic ideas.

## Supporting information

Details of datasets and Supplemental Table 1,2 and Fig 1,2

## Acknowledgment

SS supported by NPDF, SERB-DST, India, File ID: PDF/2021/000585. PC and AB acknowledge NBRC Flagship program, DBT, India, Award ID: BT/MED-III/NBRC/Flagship/Flagship2019. DR was supported by Ramalingaswami Fellowship, Department of Biotechnology (DBT), India, Award ID: BT/RLF/Re-entry/07/2014 and Department of Science and Technology (DST), India, Award ID: SR/CSRI/21/2016. The Ministry of Youth A airs and Sports, India supported AB, Award ID: F.NO.K-15015/42/2018/SP-V.

