## Supplementary material for "Neuromolecular interactions guiding homeostatic mechanisms underlying healthy ageing: A view from computational microscope": Details of datasets and Supplemental Table 1,2 and Fig 1,2

March 23, 2023

#### 1 Details of datasets used

##### Berlin data

###### Subjects

The study included 36 healthy subjects (22 females, 14 males). The written consent from the participants are taken care by [Schirner et al., 2015]. The earlier study [Schirner et al., 2015] was approved by ethics committee of the Charité University Berlin, and all experiments were performed in compliance with the relevant laws and institutional guidelines. We divided 41 subjects into two age groups comprising 22 young participants ranged in age range 18-33 years (mean age=25.68 ± 4 years, 13 female) and 16 elderly participants of age range 55-80 years (mean age=65.06 ± 7.39 years, 11 female).

###### Data Acquisition

T1 structural magnetic resonance images (MRI) and diffusion-weighted images (DWI) were acquired at Berlin Center for Advanced Imaging, Charité University Medicine, Berlin, Germany. MRI was performed on a 3T Siemens Trim Trio scanner and a 12 channel Siemens head coil (voxel size). Structural (T1-weighted high-resolution three-dimensional MPRAGE sequence; TR=1900ms, TE=2.52ms, TI=900ms, flip angle=9°, field of view (FOV)=256mm×256mm×192mm, 256×256×192 matrix, 1.0mm isotropic voxel resolution), diffusion-weighted (T2-weighted sequence; TR=7500ms, TE=86ms, FOV=192mm×192mm, 96×96 matrix, 61 slices, 2.3mm isotropic voxel resolution, 64 diffusion directions), and fMRI data (2-dimensional T2-weighted gradient echo planar imaging blood oxygen level-dependent contrast sequence; TR=1940ms, TE=30ms, flip angle=78°, FOV=192 × 192mm$^2$, 3 × 3mm$^2$ voxel resolution, 3mm slice thickness, 64 × 64 matrix, 33 slices, 0.51ms echo spacing, 668 TRs, 7 initial images were acquired and discarded to allow magnetization to reach equilibrium; eyes-closed resting-state) were acquired on a 12-channel Siemens 3 Tesla Trio MRI scanner at the Berlin Center for Advanced Neuroimaging, Berlin, Germany.

###### Structural connectivity

The structural connectivity (SC) for each subject and age is generated using the pipeline described by Schiner et. al. [Schirner et al., 2015]. In the pipeline, high-resolution T1 anatomical images were used to create segmentation and parcellation of cortical and sub-cortical gray matter, white matter segments. The main pre-processing steps for T1 anatomical images involved skull stripping, removal of non-brain tissue, brain mask generation, cortical reconstruction, motion correction, intensity normalization, WM, and subcortical segmentation, cortical tessellation generating GMWM and GM-pia interface surface-triangulation and probabilistic atlas-based cortical and subcortical parcellation. Cortical grey matter parcellation of 34 region of interest (ROI) in each hemisphere was undertaken following Desikan-Killiany parcellation [Desikan et al., 2006]. The connection strength (a value ranging from 0 to 1) between each pair of ROIs was estimated by probabilistic tractography algorithm. SC matrices were generated from each subject's MRI data and then summed element-wise to obtain an averaged SC matrix. The pre-processing steps for the diffusion MRI data were eddy current and motion correction with re-orientation of b-vectors (b-zero image was linearly registered to the subject's anatomical T1-weighted image). Tractography was constrained by seed, target, and stop masks. The fiber length was represented in millimeters. The 68 ROIs or nodes had no self-connection loops meaning that the diagonal values of the SC matrix are all zero.

##### Resting state functional connectivity

The participants were subjected to a functional MRI scan, when eyes-closed awake resting-state data were acquired. The resting-state BOLD activity was recorded for 22 minutes (TR=2 sec) using a 3T Siemens Trim Trio scanner, and a 12 channel Siemens head coil (voxel size). The BOLD signal was computed for 68 ROIs, parcellated by Desikan-Killiany atlas [Desikan et al., 2006] and z-transformed. Pearson correlation coefficient, between each pair of region, defines resting-state FC matrix.

#### NKI/Rockland data set:

We tested our hypothesis on another human neuroimaging dataset obtained from Nathan Kline Institute (NKI)/Rockland sample (available in the UCLA Multimodal Connectivity Database (UMCD)[Brown et al., 2012].

##### Structural connectivity

Diffusion Tensor Imaging (DTI) based structural connectivity matrices and rsFC/Pearson correlation matrix of 75 healthy participants were obtained from the Nathan Kline Institute (NKI), Rockland sample [Brown et al., 2012]. The participants consisted of 30 males and 35 females, with a total age range of 19-85 (mean ± std). We separate 40 participants into two age groups (48 young and 27 old) and averaged their SCs over individual subject for our analysis (see Table 1). Cortical gray matter was parcellated using the Craddock 200 atlas [Craddock et al., 2012]. Elements of the SC represented the number of white matter tracts between gray matter parcels. The reader may also refer to the original work [Brown et al., 2012] for details regarding structural and functional connectome pre-processing. Prior to running the models on the NKI SC, the SC was scaled down by dividing every element by the maximum value found in the original SC consisting of white matter tract numbers between regions. Diffusion tensors were estimated using Diffusion Toolkit (http://trackvis.org/blog/tag/diffusion-toolkit) and tractography was run using the fiber assignment by continuous tracking algorithm [Mori and Van Zijl, 2002]. For each ROI, all fibers were counted that intersected at least one voxel in the source ROI and at least one voxel in any target ROI using custom code, (http://ccn.ucla.edu/wiki/index.php/UCLA_Multimodal_Connectivity_Package). Thus, 188×188 structural connectivity was obtained.

##### Resting-state functional connectivity

Resting state fMRI data was pre-processed using the pipeline described in [Brown et al., 2012]. After marking flagged TRs, the mean time series for each ROI was calculated and then correlated with all remaining ROI time series (excluding flagged TRs) to derive a 188×188 resting-state functional connectivity (rsFC) matrix.

#### Cam-CAN data set

##### Subjects

In our study, total 69 healthy participants were included from the Cambridge Centre for Ageing and Neuroscience (Cam-CAN) cohort http://www.mrc-cbu.cam.ac.uk/datasets/camcan/http://www.mrc-cbu.cam.ac.uk/datasets/camcan/ [Shafto et al., Taylor et al., 2017].

##### Data acquisition

Resting state MRI (T1-weighted image), diffusion weighted MRI, and functional MRI were performed using a 3T Siemens TIM Trio scanner with a 32-channel head-coil at Medical Research Council (UK) Cognition and Brain Sciences Unit (MRC-CBSU) in Cambridge, UK.

High-resolution 3D T1-weighted data was collected with a magnetization prepared rapid gradient echo (MPRAGE) sequence using Generalized Autocalibrating Partially Parallel Acquisition (GRAPPA) with acceleration factor of 2 and other parameters were: repetition time (TR)=2,250 ms, echo time (TE)=2.99 ms, flip angle: 9°, field of view (FOV)=256 × 240 × 192 mm; voxel size = 1 $mm^3$ isotropic, inversion time (TI)=900 ms, acquisition time (TA)=4:32min [Shafto et al., 2014].

Diffusion data were acquired in a twice-refocused spin echo sequence with TR of 9100 ms, TE of 105 ms, FOV of 192 × 192 mm, isotropic voxel size of 2 $mm^3$, 66 axial slices using 30 directions with b = 1000 $s/mm^2$, 30 directions with b = 2000 $s/mm^2$, and three b = 0 images with single average [Shafto et al., 2014]. Resting state eye closed fMRI was collected with EPI sequence with 1970 ms TR, 30ms TE, 78° flip angle, 192 × 192 mm FOV, 3 × 3 × 4.44 $mm^3$ voxel size, 32 slices of thickness 3.7 mm [Shafto et al., 2014].

##### DTI processing

Diffusion MRI data were processed locally using own pre-processing pipeline by using MRtrix3 ([Tournier et al., 2019], https://github.com/MRtrix3/mrtrix3https://github.com/MRtrix3/mrtrix3), FSL http://www.fmrib.ox.ac.uk/fsl http://www.fmrib.ox.ac.uk/fsl, and ANTS http://stnava.github.io/ANTs/, http://stnava.github.io/ANTs/. Main preprocessing steps include denoising (MRtrix command 'dwidenoise', [Veraart et al., 2016]), Gibb's ringing artefacts removal (MRtrix command 'mrdegibbs', [Kellner et al., 2016]), motion and eddy current corrections (MRtrix command 'dwifslpreproc', [Andersson et al., 2003]), biasness corrections (MRtrix command 'dwibiascorrecT using ANTs, [Tustison et al., 2010]). Then a brain mask in DTI space was calculated for each subject using 'beT command ( in FSL) on the preprocessed image. To find the orientation of the fiber(s) (fiber orientation distribution, FOD) in each voxel, we used multi-shell multi-tissue constrained spherical deconvolution (MSMT-CSD; MRtrix command 'dwi2response' and 'dwi2fod', [Dhollander et al., 2018]). After that, a global intensity normalization has been performed to make the fiber orientation distributions comparable between subjects. We used Anatomically Constrained Tractography (ACT, MRtrix command '5ttgen', [Smith et al., 2012]) to generate a tissue-segmented image. In order to use ACT, we first preprocess the T1-weighted image using MRtrix, then we co-registered that image to the DWI using both MRtrix and FSL. We created an mask of the gray-matter/white-matter-boundary, which was useful for streamline seeding. Finally, probabilistic tractography (MRtrix command 'tckgen') has been used to generate 20 million tracks. The tractogram was filtered (SIFT2 approach, MRtrix command 'tcksift2') further to a find subset of streamlines such that the streamlines densities were more close to fibre densities [Smith et al., 2015]. Each steps were visually assessed and edited by research personnel.

##### T1 processing

Freesurefer http://surfer.nmr.mgh.harvard.eduhttp://surfer.nmr.mgh.harvard.edu [Dale et al., 1999] 'recon-all' used to reconstruct a two-dimensional cortical surface from a three-dimensional volume acquired from T1-weighted image. Recon-all steps were skull stripping from the anatomical image, estimation of interface between the white matter and grey matte, generation of white and pial surfaces. Cortical parcellation was performed using the Desikan-Killiany atlas [Desikan et al., 2006], with 68 ROIs.

##### Structural connectivity

A Desikan–Killiany atlas based whole-brain connectome was generated for each subjects by computing the fiber density between each pair of ROIs (MRtrix command 'tck2connectome' with option 'scale_invnodevol') to count how many streamlines from one region reach every other regions.

##### fMRI preprocessing

Functional MRI (fMRI) images for resting state has been preprocessed using CONN toolbox [?] https://web.conn-toolbox.org/https://web. conn-toolbox.org/, a Matlab/SPM-based software. Data were preprocessed using a default preprocessing pipeline of CONN. The preprocessing methods were unwrapping using field-map images, realignment to correct for motion, slice timing correction, segmentation, normalisation to the MNI template, outlier rejection and functional smoothing. Spatial smoothing was performed using a Gaussian kernel with full width of 6.0 mm. Denoising was used to remove signal changes related to white matter, cerebrospinal fluid, motion, breathing and cardiac pulsations. Finally temporal band-pass filtering at (0.04-0.9 Hz) and linear detrending were performed. For region of interest (ROI) analysis, mean regional BOLD time series were estimated in 68 parcellated brain areas of Desikan–Killiany atlas [Desikan et al., 2006].

##### Resting state functional connectivity

The same participants were subjected to a functional MRI (fMRI) scan during which their eyes-closed awake resting-state data were acquired. The resting-state BOLD activity was recorded for 22 minutes (TR=2 sec). The BOLD activity was then down-sampled to fit the 68 ROIs. Aggregated BOLD time series of each region were z-transformed. Next, pairwise Pearson correlation coefficient were computed to obtain $68 \times 68$ rsFC matrix for individual subjects.

Table 1: Demographic data across two different stages of adulthood in men and women.

| Data set | Group | Young<br>Number/mean±sd | Old<br>Number/mean±sd |
|---|---|---|---|
| Cam-CAN | | Age range 18-34 | Age range 60-85 |
| | Female | 18 / 26.28 ± 4.43 | 18 / 68.1 ± 6.67 |
| | Male | 14 / 25.57 ± 5.17 | 19 / 70.16 ± 8.08 |
| | Total | 32 / 25.97 ± 4.7 | 37 / 69.14 ± 7.41 |
| Berlin | | Age range 18-33 | Age range 55-80 |
| | Female | 13 / 26 ± 4.73 | 11 / 64.13 ± 7.74 |
| | Male | 12 / 25.33 ± 3.2 | 3 / 58.75 ± 10.4 |
| | Total | 25 / 25.68 ± 4 | 14 / 62.3 ± 8.71 |
| NKI | | Age range 18-30 | Age range 60-85 |
| | Female | 19 / 22.58 ± 2.85 | 16 / 69.69 ± 7.92 |
| | Male | 29 / 22.72 ± 2.81 | 11 / 70.27 ± 9.3 |
| | Total | 39 / 23.08 ± 3.12 | 27 / 70.29 ± 7.62 |

### 2 Results

#### Network property analysis

We validate our observations on three data sets, The results are shown in the Fig. S4.

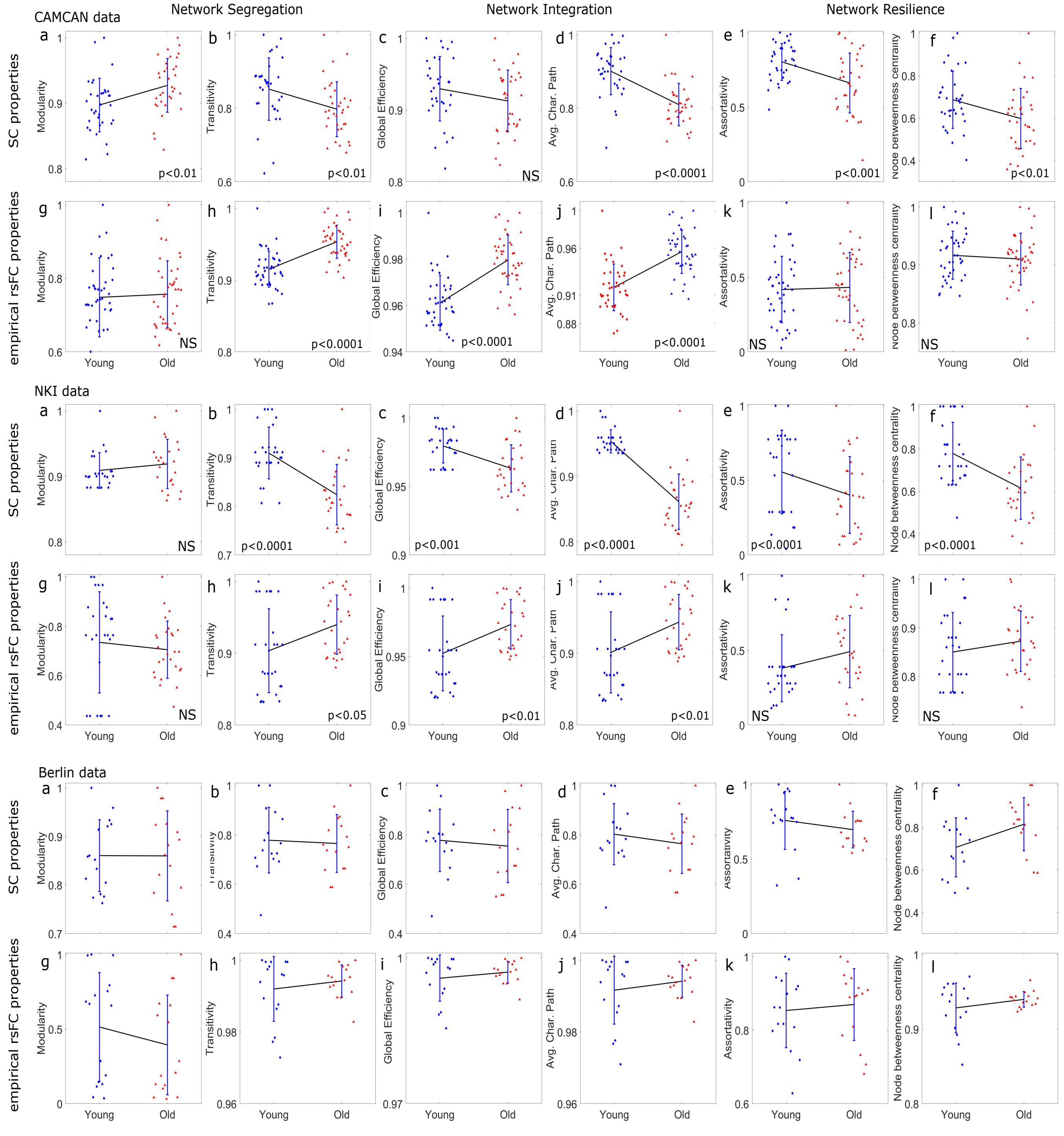


Figure 1: Changes in SC and functional network properties in the three data sets. To show pattern of the shift between two means of the two groups, we joint then by a line, significance alteration (or unchanged) network properties are determined by two-sample t-test.

#### Multiscale Dynamic Mean Field (MDMF) model

The whole-brain dynamics is described by a set of coupled nonlinear stochastic differential equations, the current-based MDMF model [Naskar et al., 2021] as,

$$
\begin{aligned}
[!h] I_i^{(E)} &= w_E I_0 + w_+ J_{NMDA} S_i^{(E)} + G J_{NMDA} \sum_{j=1}^{N} C_{ij} S_j^{(E)} - J_i S_i^{(I)} \\
I_i^{(I)} &= w_I I_0 + J_{NMDA} S_i^{(E)} - c\, S_i^{(I)} \\
r_i^{(E)} &= \frac{a_E I_i^{(E)} - b_E}{1 - exp^{-d_E\left(a_E I_i^{(E)} - b_E\right)}} \\
r_i^{(I)} &= \frac{a_I I_i^{(I)} - b_I}{1 - exp^{-d_I\left(a_I I_i^{(I)} - b_I\right)}} \\
\frac{dS_i^{(E)}}{dt} &= -\beta^E S_i^{(E)} + \alpha^E \left(1 - S_i^{(E)}\right) T_{glu}\, r_i^{(E)} + \sigma \nu_i(t) \\
\frac{dS_i^{(I)}}{dt} &= -\beta^I S_i^{(I)} + \alpha^I \left(1 - S_i^{(I)}\right) T_{gaba}\, r_i^{(I)} + \sigma \nu_i(t) \qquad (1)
\end{aligned}
$$

where, the subscripts $i, j$ indicate brain areas, and superscripts $E, I$ represent excitatory or inhibitory populations, respectively. Total number of considered brain areas is $N$, where its value is different for different parcellation choice, e.g., $N = 68$ and 188, respectively for Desikan-Killiany parcellation [Desikan et al., 2006] and Craddock 200 atlas [Craddock et al., 2012]. $S_i^{(E,I)}(t)$ denotes the average excitatory or inhibitory synaptic gating variables at $i^{th}$ local brain area. The time-dependent gating variables are drawn by averaging the fraction of open channels of neurons. The stochasticity is introduced by adding uncorrelated white Gaussian noise $\nu_i$ within the equation of two gating variables with intensity $\sigma$ for each brain region. Further details of the model formulation and descriptions are available in the article by A. Naskar et al. [Naskar et al., 2021].

$J_i$ denotes the local synaptic coupling strength from inhibitory to excitatory. The temporal dynamics of the inhibitory feedback are given by,

$$
\frac{dJ_i}{dt} = \gamma r_i^I (r_i^E - \rho) \qquad (2)
$$

At the mean-field level, the biological complexity involved in the balance of dynamics between excitatory and inhibitory fields can be captured grossly using the mathematical implementation of the inhibitory plasticity rule [Hellyer et al., 2016]. An inhibitory plasticity rule represents changes in $J_i(t)$ (synaptic weight) to ensure that the inhibitory current clamps to an excitatory population, maintaining homeostasis. Homeostasis is achieved with $J_i(t)$ dynamics such that the firing rate of the excitatory population is maintained at the target firing rate $\rho = 3Hz$, and $\gamma$ is the learning rate in $sec$. The chosen target firing rate is the firing rate observed when the inhibitory and excitatory currents are matched.

Table 2: Default parameter values, and descriptions shown with references.

| Parameter | Value/Unit | Description | Ref. |
|---|---|---|---|
| $I_0$ | $0.382\ nA$ | overall effective external input current | [Deco et al., 2014] |
| $w_+$ | 1.4 | local excitatory recurrence | [Deco et al., 2014] |
| $J_{NMDA}$ | $0.15\ nA$ | excitatory synaptic coupling strength | [Deco et al., 2014] |
| $\sigma$ | 0.001 | noise intensity | [Naskar et al., 2021] |
| $w_E$ | 1 | scaling parameter for excitatory populations | [Deco et al., 2014] |
| $w_I$ | 0.7 | scaling parameter for inhibitory populations | [Deco et al., 2014] |
| $\alpha_E$ | $0.072\ mMol^{-1}$ | forward rate constants of excitatory pools | [Destexhe et al., 1994] |
| $\alpha_I$ | $0.53\ mMol^{-1}$ | forward rate constants of inhibitory pools | [Destexhe et al., 1994] |
| $\beta_E$ | $0.0066\ ms^{-1}$ | backward rate constants of excitatory pools | [Destexhe et al., 1994] |
| $\beta_I$ | $0.18\ ms^{-1}$ | backward rate constants of inhibitory pools | [Destexhe et al., 1994] |
| $a_E$ | $310\ nC^{-1}$ | parameter for input-output function | [Deco et al., 2014] |
| $a_I$ | $615\ nC^{-1}$ | parameter for input-output function | [Deco et al., 2014] |
| $b_E$ | $125\ Hz$ | parameter for input-output function | [Deco et al., 2014] |
| $b_I$ | $177\ Hz$ | parameter for input-output function | [Deco et al., 2014] |
| $d_E$ | $0.16\ s$ | parameter for input-output function | [Deco et al., 2014] |
| $d_I$ | $0.087\ s$ | parameter for input-output function | [Deco et al., 2014] |
| $\gamma$ | $1\ C$ | learning rate | [Naskar et al., 2021] |
| $\rho$ | $3\ Hz$ | target firing rate | [Deco et al., 2014] |
| $G$ | 0.5 | global coupling strength | |
| $c$ | $1\ nA$ | local inhibitory to inhibitory strength | |
| $T_{glu}$ | 0.1-15 $mMol$ | glutamate concentration | [Naskar et al., 2021] |
| $T_{gaba}$ | 0.1-15 $mMol$ | GABA concentration | [Naskar et al., 2021] |

#### Model simulation and GGC estimation

Figure 2 describes the computational microscope and four steps to estimate synthetic concentrations of the two primary neurotransmitters.

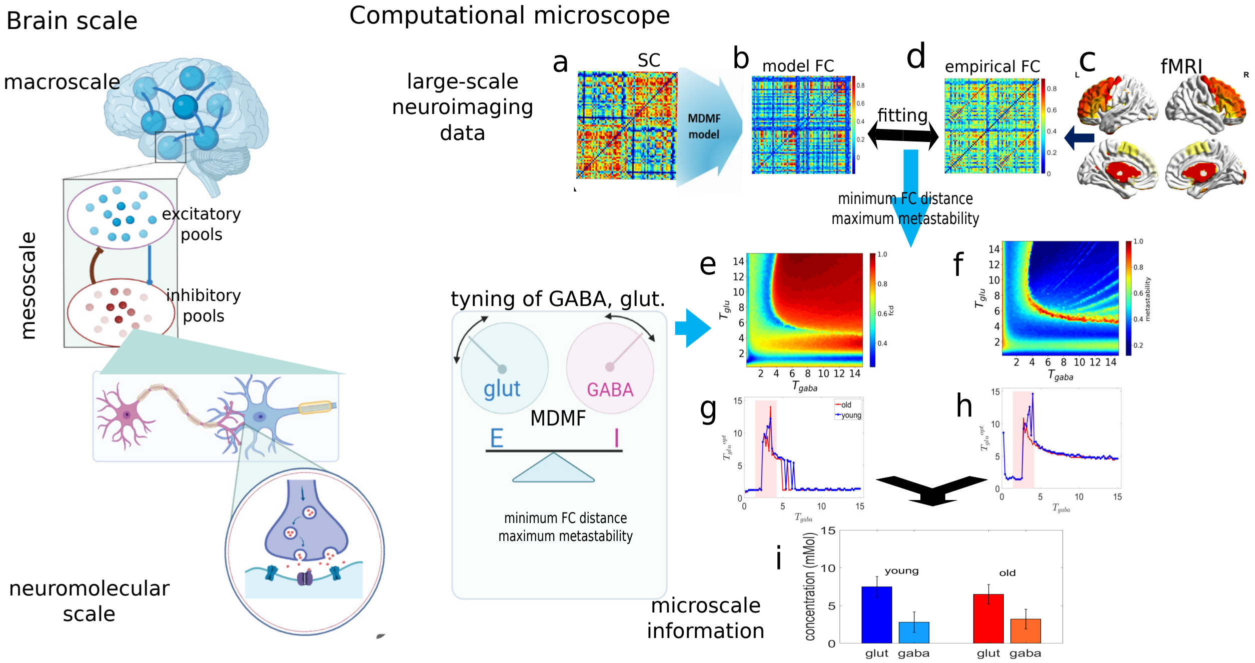


Figure 2: **Schematic representation for estimating synthetic GABA and glutamate concentrations using computational microscope.** Four steps of the model simulation and synthetic result generation are shown here. (a-d) Step 1, we compare the model FC from spatiotemporal patterns to those observed in the empirical FC obtained from fMRI BOLD activity at rest. (e, f) In step 2, FCD and metastability are stored for each pair of GABA-glutamate concentrations. (g, h) Step 3, we estimate optimal glutamate for different values of GABA imposing the two optimal conditions. (i) Step 4, an optimal GABA is estimated from the result obtained in the previous step. Thus, an optimal set of GGC is estimated, based on minimum FCD and maximum metastability.

Step 1, in Figs. 2(a-d), involves synthetic generation of neural activity and fitting method that allows us to project the model generated low frequency $(0.004 - 0.1\text{Hz})$ BOLD signals and their spatiotemporal harmony, particularly fluctuations in the cortical coherence. The spontaneous brain activity at rest is captured in form of the empirical BOLD signals derived from fMRI data. The two BOLD signals (empirical and simulated) depicts hemodynamics responses of different brain areas, where pair-wise spatial correlation is the weights in functional connectivity (FC) matrix. Detailed steps in fist stage analysis are: Fig. 2(a) We use anatomical connectivity derived by diffusion tensor imaging from the subjects of different ages. Fig. 2(b) A mean-field neuronal model is put on top of the structural connectivity matrix. Thus, the nodal dynamics is governed by the MDMF model [Naskar et al., 2021] and spatially connected via SC matrix. The structural connectivity describes the inter-areal interaction strength to exchange excitability across the cortical regions. We generate a model simulated neural activity, firing rate of excitatory/inhibitory populations of individual regions. The Balloon-Windkessel algorithm [Friston et al., 2003] then estimates the hemodynamic activity at rest. A Pearson correlation is derived from the estimated hemodynamic activity between each pair of brain regions. Thus, we approximate the functional connectivity (FC), which we mentioned as the model based or simulated FC. Figure 2(c) shows an exemplary resting-state fMRI activation map, which is used to extract empirical resting-state BOLD activity of individual subjects using CONN toolbox. In Fig, 2(d), we plot the empirical rsFC from the BOLD activity, which is used in fitting against resting-state model FC.

In Step 2, the fitting measures euclidean distance between empirical and model FCs, called functional connectivity distance (FCD), shown in Fig. 2(e). Simulteneously, the fluctuation in global coherence is measured by metastability, when tuning the two intrinsic parameters ($T_{glu}$ and $T_{gaba}$), see Fig. 2(f). We have varied the two local parameters, $T_{glu}$ and $T_{gaba}$ homogeneously across the whole brain and store the two measures for each pair of parameter values in the two-parameter phase diagrams, see Figs. 2(e) and 2(f).

In Step 3, see Figs. 2(g,h), we scan along x-axis, i.e., across GABA values, and estimate optimal glutamate concentration ($T_{glu}$) in $mMol$ imposing two optimal conditions, minimum FC distance (i.e., maximum correlation between the model generated FC and empirical rsFC), and maximum metastability, which decides optimal brain functionality. We average over the estimated glutamate values obtained from the two measures from Figs. 2(g,h).

Finally, in Step 4, GABA concentration is estimated for the homeostasis range [Naskar et al., 2021], indicated by shaded boxes in Figs 2(g,h) under the same optimal conditions. Thus subject wise optimal parameter set of $T_{gaba}$ and $T_{glu}$, is predicted investigating whole-brain neuroimage data (MRI, fMRI), shown using a representative result in Fig. 2(i).

Further, we monitor GABA-glutamate concentrations (GGC) and their ratio (GGR) on a very slow time scale (18-84 years) at the level of single subject. We separate healthy subjects into two age groups, young and old. Then, a possible alteration (or invariance) pattern in the two metabolites is captured over lifespan, which helped to draw plausible conclusion, see lower box in Fig. 2.

#### Model-based rsFC

Here we discussed how we estimate model generated functional connectivity matrix, which will be used to check the model's predictability under certain assumptions. The simulated neural fluctuations were given by the gating variable $S_i^{(E)}(t)$ and firing rate $r_i^{(E)}(t)$ for $i^{th}$ brain area fluctuates around a fixed value. The neural activity from the MDMF model was converted to the BOLD activity using a hemodynamic model [Friston et al., 2003]. The method was adopted from previously published works [Cabral et al., 2011]. The simulated BOLD activity was filtered by a band pass filter followed by Fisher z-transform converting into normal distribution. Pearson correlation between all pair of regions generates the model FC.
